# The effects of breastmilk-derived osteopontin on the intestinal intraepithelial lymphocyte compartment

**DOI:** 10.1101/2025.05.29.656804

**Authors:** Kathleen G. McClanahan, Jayden C. Capella, Jennifer A. Gaddy, Danyvid Olivares-Villagómez

## Abstract

Osteopontin is a protein with many physiological roles and is widely expressed by many cell types, tissues, and bodily fluids, including breastmilk. The functions of breastmilk osteopontin are not clearly defined, however, it is known to impact intestinal and brain development in infants. Although it is known that endogenous osteopontin influences the survival of intestinal intraepithelial lymphocytes (IEL)^2^, the impact of breastmilk osteopontin on developing intestinal immune cells remains unclear. In this report, mouse models lacking expression of osteopontin were used to demonstrate that milk-derived osteopontin is important for the development of IELs, with observed effects in both juvenile and adult mice. These changes are most prevalent in IELs expressing CD8αα: however, the impact of these alterations is unclear, as mice with disrupted IEL compartments are not more susceptible to DSS-induced colitis or infection by *Citrobacter rodentium*.

## Introduction

Infancy is a time of great physiological change and stress. This period is characterized by rapid growth and development, corresponding with exposure to numerous environmental factors. One of the main mechanisms through which the infant is exposed to the environment is through the gastrointestinal tract, which through its large surface area allows contact between the infant and numerous nutrients, microbes, and foreign factors. In the infant, intestinal and immune development are not yet complete, resulting in a period of heightened susceptibility to pathogens and other stimuli. The main mechanism through which infants are protected during this period is breastfeeding. Breastmilk is a complex biological fluid composed of thousands of distinct compounds, the most prominent of which support infant nutrition, while others have bioactive potential. Bioactive proteins including IgA, lactoferrin, and lysozyme play roles in protecting the infant from potential pathogens (1)(2)(3)(4). Other components, including human milk oligosaccharides (HMOs), act as prebiotic factors, promoting the development of a robust intestinal microbiota and preventing the colonization of pathogenic organisms(5).

One abundant bioactive protein present in breastmilk is osteopontin, a highly phosphorylated glycoprotein with numerous physiological roles. Originally isolated from rat bone matrix(6), osteopontin has been shown to impact calcification processes(7), adhesion and migration of cells(8)(9), and immune cell function(10)(11), among others. In addition to being found in most tissues, osteopontin is also present in many bodily fluids, including breastmilk(12).

Osteopontin plays important roles in the development of the infant. Mouse models have demonstrated the importance of milk-derived osteopontin in the development of the intestinal epithelium(13)(14), as well as in brain myelination(15). Studies in rhesus macaques demonstrated that supplementation of formula with osteopontin led to an intestinal epithelial cell transcriptome more similar to that of breastfed infants(16). Trials of osteopontin supplementation in human infants have been promising, with infants fed osteopontin-supplemented formula showing reduced rates of fever, as well as a reduction in inflammatory cytokines such as TNF-α and IL-6(17). In Europe, a bovine osteopontin supplement, known as Lacprodan® OPN-10, has been approved for use in infant formula and has shown a robust safety profile(18)(19). Given the potential for commercially available osteopontin-supplemented formulas in the near future, it is of critical importance to understand the effects of osteopontin on infant development. One known target of osteopontin in the adult intestine is the intraepithelial lymphocyte (IEL) compartment. IEL are immune cells that reside between intestinal epithelial cells, interspersing at a ratio of ∼1 IEL for every 10-20 epithelial cells in the small intestine and ∼1 IEL for every 40 epithelial cells in the colon(20)(21). IEL play critical roles in intestinal homeostasis, including immune tolerance and intestinal defense(22)(23)(24). IEL come in many subtypes; many of which straddle the bridge between innate and adaptive immunity. Many IEL express either TCRαβ or TCRγδ; however, most IEL populations are limited in antigen recognition and/or respond to antigen through mechanisms other than canonical TCR signaling(25, 26). The most prevalent IEL subsets include TCRβ^+^CD4^+^ and TCRβ^+^CD8α^+^ IELs, as well TCRγδ^+^ IELs in the small intestine. Of note, many CD8α^+^ IEL do not express the canonical CD8αβ co-receptor and instead express an alternate CD8αα form. Many IELs develop during early life(27), leading us to hypothesize that milk-derived osteopontin may play a role in their development or maturation.

We found that osteopontin derived from maternal breastmilk plays a role in IEL development, as mice that did not receive milk osteopontin displayed alterations in the IEL compartment both as juveniles (3w) and adults (8w). However, the impact of these alterations remains unclear, as mice with altered IEL compartments were not more susceptible to the tested models of intestinal inflammation.

## Materials and Methods

### Mice

All experiments and animal procedures were conducted according to the NIH guidelines for the care and use of laboratory animals, under a protocol approved by the Institutional Animal Care and Use Committee at Vanderbilt University. C57BL6/J and osteopontin KO (B6.129S6(Cg)-*Spp1^tm1Blh^*/J) mice were originally purchased from The Jackson Laboratory (000664 and 004936, respectively) and have been maintained in our colony for several years. Heterozygous mice were generated by crossing *Spp1*^-/-^ with WT C57BL6/J mice. F1 males (*Spp1*^+/-^) and females (*Spp1*^+/-^) were then crossed to generate *Spp1*^+/+^, *Spp1*^+/-^, and *Spp1*^-/-^ littermate offspring. Littermate *Spp1*^+/-^ and *Spp1*^-/-^ females were bred with *Spp1*^+/+^ males to generate pups for experiments. Females were harem bred (3-4 females and 1 male per cage) and maintained in the same breeding cage until external signs of pregnancy, at which point the pregnant female was removed to a fresh cage. Mice were maintained in a temperature (21-22°C) controlled room under a 12-hour light/dark cycle. Water and food were supplied *ad libitum*. Offspring were weaned at postnatal day (PND)^3^ 21. No significant sex differences were found in experiments in this study, and male and female mice were used for all experiments except for those related to milk production.

### Cross-fostering of mouse pups

Littermate *Spp1*^+/-^ and *Spp1*^-/-^ dams were timed mated with *Spp1*^-/-^ sires. Briefly, *Spp1*^+/-^ and *Spp1*^-/-^ dams were maintained in exclusively female cages to allow synchronization of the estrous cycle. Female mice were then set to breed with singly housed *Spp1*^-/-^ sires that had been isolated for >5 days to allow for maximum sperm count. Females were left to breed for 1-4 days before being separated from the male. Females were observed for external signs of pregnancy, at which point they were separated into a clean cage. At PND0 or PND1, dams of time-matched litters were transferred between cages (*Spp1*^-/-^ dam to pups from *Spp1*^+/-^ dam and vice versa). Mice were monitored for signs of pup acceptance and feeding. Pups were left with the foster dam until use in an experiment or until weaning at PND21. If exact time-matched litters were not available, pups were occasionally switched on PND2 or PND3. In this case, the younger litter was always given to the *Spp1*^-/-^ dam to minimize milk osteopontin exposure.

### Milking of mouse dams

Dams were separated from litter into a clean cage 2 hours prior to milking. After 2 hours had elapsed, dams were injected i.p. with 2 IU of oxytocin (Sigma Aldrich, O3251) and placed back into the cage for 3-5 minutes while milk letdown was initiated. Dams were securely scruffed and milk was collected from at least 2 nipples using an adapted human breast pump (similar to that reported by(28)). Milk was stored at −70°C until analysis.

### Flush of mouse pup intestines

PND2, PND5, PND7, or PND21 pups were euthanized using CO_2_, followed by decapitation. The small intestines and colons (minus cecum) were removed and stored on ice in tubes containing 1 ml of PBS with cOmplete^TM^, EDTA-free Protease Inhibitor Cocktail (Roche, 04693132001, 1 tablet per 10 mL PBS). Intestines and their contents were homogenized using a handheld homogenizer (Fisherbrand) then centrifuged to pellet debris. Supernatant was transferred to clean tubes and stored at −70°C until analysis. PND28, PND35, and PND42 mice were similarly processed, except only sections of the small intestine (combined proximal, medial, and distal) were processed.

### Determination of osteopontin protein concentration

Previously frozen breastmilk and intestinal flush samples were brought to room temperature on the counter. Intestinal flush samples were centrifuged for 10 minutes at 10,000xg, 4°C prior to analysis. Samples were analyzed using Osteopontin (OPN/SPP1) Mouse ELISA Kit (Invitrogen, EMSPP1), according to kit instructions. Intestinal samples were not diluted prior to analysis, milk samples were diluted 1:1,000,000 (1:1000 dilution followed by 1:1000 dilution).

### Lymphocyte Isolation

Spleen and mesenteric lymph nodes (MLN)^4^ lymphocytes were isolated by conventional means(29). Briefly, cells were manually dispersed through a 70 mM cell strainer (ThermoFisher, 22-363-548) and ACK lysing buffer (KD Medical, RGF-3015) was applied to cell pellets to lyse red blood cells. Cells were recovered into cold RPMI (Gibco, 11835-030) supplemented with 10% fetal bovine serum (R&D Systems, S11150H), 2 mM L-glutamine (Corning, 25-005), 1x penicillin-streptomycin (Corning, 30-002), 10 mM HEPES (Corning, 25- 060), and 25 mM 2-mercaptoethanol (Gibco, 21985-023). IEL cells were isolated by mechanical disruption as previously reported.(30) Briefly, after flushing the intestinal contents with cold HBSS, cutting longitudinally, and removing excess mucus, the intestines were cut into small pieces (∼1 cm) and shaken for 45 minutes at 37°C in HBSS supplemented with 5% fetal bovine serum and 2mM EDTA (KD Medical, RGF-3130). Supernatants were recovered and cells isolated using a discontinuous 40%/70% Percoll (Cytiva, 17089101) gradient.

### Cell Staining

Surface cell staining was performed following conventional techniques. For intracellular cytokine staining, cells were stimulated with Cell Activation Cocktail without Brefeldin A (BioLegend, 423302) in the presence of Golgi Stop (BD Biosciences, 51-2092KZ) for 4 hours prior to staining. Extracellular markers were stained, cells were fixed briefly with 2% paraformaldehyde (Electron Microscopy Sciences, 15710-S), followed by permeabilization and intracellular staining using the BD Cytofix/Cytoperm kit (BD Biosciences, 554714) according to manufacturer’s instructions. All stained samples were acquired using BD FACS Canto II or BD LSRFortessa (BD Biosciences). Data were analyzed using FlowJo software (FlowJo LLC). Antibodies used can be found in Table I. Gating strategies can be found in Figure S1.

### DSS-Induced Colitis

Mice were supplied with 3% DSS (Thermo Scientific, J63606.22) in the drinking water for 5 days before being switched back to sterile water. Mice were weighed daily and monitored for signs of disease, including rectal inflammation, diarrhea, and wasting. For the recovery model, mice were exposed to 4% DSS for 5 days before returning to sterile water.

#### Citrobacter rodentium infection

Mice were infected with 1×10^9^ CFU of *Citrobacter rodentium* (ATCC, 51459) by oral gavage. Mice were weighed daily and monitored for signs of disease, including rectal inflammation, diarrhea, and wasting, until harvest at 14 days post-infection.

### Statistical analysis

Data were analyzed using Prism 9 (GraphPad, San Diego, CA) to determine significant (p < 0.05) differences between groups. Comparisons between two groups were made by Student’s t test. Data are presented as mean ± standard error of the mean.

## Results

To determine osteopontin breastmilk concentration in our mouse models, we analyzed milk from *Spp1*^+/+^, *Spp1*^+/-^, and *Spp1*^-/-^ dams from our colony at 5 days post-partum (PND5) (Figure 1A). As expected, milk from *Spp1*^-/-^ dams had no detectible osteopontin. It is well-known that milk osteopontin concentration varies widely between individuals, which was reflected in our samples. On average, *Spp1*^+/-^ dams had roughly half the osteopontin concentration of *Spp1*^+/+^ dams; however, inter-group variance was large, with *Spp1*^+/+^ mice having milk osteopontin concentrations between 100 and 500 mg/mL, and *Spp1*^+/-^ mice having concentrations between 50 and 300 mg/mL.

**Figure 1:**
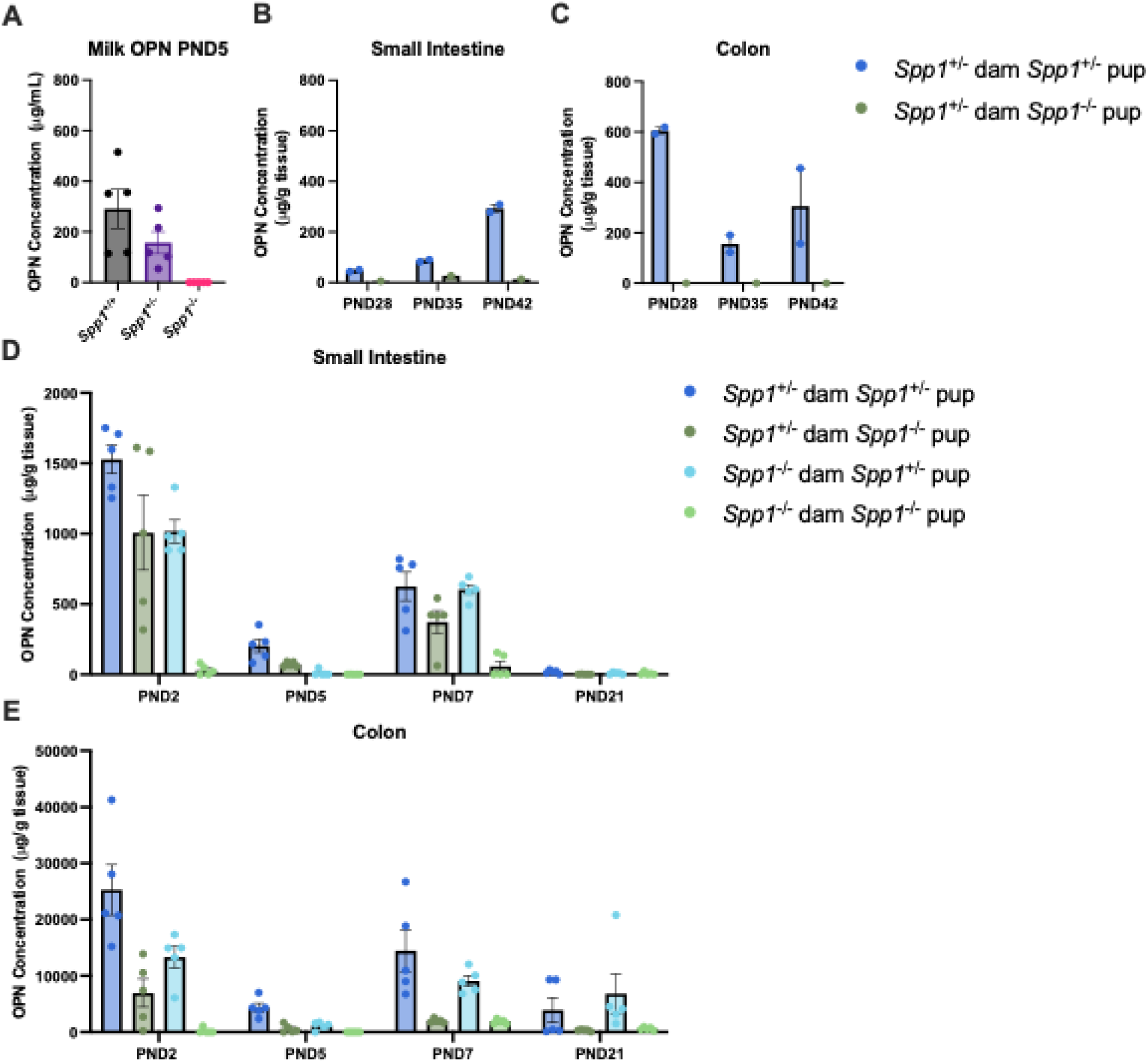
Osteopontin protein expression in breastmilk and intestines. A, milk osteopontin concentration at 5 days post-birth. 2 independent experiments, n=5. B, osteopontin concentration in the adult mouse small intestine; one independent experiment, n=2 (*Spp1*^+/-^) n=1 (*Spp1*^-/-^). C, osteopontin concentration in the adult mouse colon; 1 independent experiment, n=2 (*Spp1*^+/-^) n=1 (*Spp1*^-/-^). D, osteopontin concentration in the juvenile small intestine; 2 independent experiments, n=5. E, Osteopontin concentration in the juvenile colon. two independent experiments, n=5. Each symbol represents an individual sample.

We then investigated osteopontin concentrations in the adult intestine. Osteopontin concentration in the small intestine of *Spp1*^+/-^ mice increased over time, from 50 mg/g at PND28 to roughly 300 mg/g at PND42 (Figure 1B). An opposite trend was seen in the colon of the same mice, with the highest concentration observed at PND28 and lower concentrations observed at PND35 and PND42 (Figure 1C). We then investigated the effects of maternal and pup genotype on intestinal osteopontin by analyzing small intestinal and colon contents of *Spp1*^+/-^ and *Spp1*^-/-^ pups from *Spp1*^+/-^ (osteopontin-competent) dams crossed with *Spp1*^-/-^ sires, and *Spp1*^-/-^ (osteopontin-deficient) dams crossed with *Spp1*^+/-^ sires. We observed that osteopontin concentration in the small intestine of *Spp1*^+/-^ pups reared by osteopontin competent dams (dark green bars) peaks early in life at approximately 1500 mg OPN/g tissue on PND2, followed by a decrease in osteopontin concentration to roughly 250 mg/g at PND5 that rebounds to approximately 600 mg/g by PND7. By PND21, little to no osteopontin is detectible (Figure 1D). This pattern was also observed in the colon, though at concentrations roughly 10x of that seen in the small intestine (Figure 1E). As expected, no osteopontin was seen in the intestines of *Spp1*^-/-^ pups from *Spp1*^-/-^ dams at any time point (light green bars). Of note, osteopontin was readily detectable in the intestines of *Spp1*^-/-^ pups reared by *Spp1*^+/-^ dams, indicating the presence of breastmilk derived osteopontin in the lumen of these mice (dark green bars). Osteopontin was also detected in *Spp1*^+/-^ pups reared by osteopontin-deficient dams (light blue bars), indicating a possible intestinal mechanism to compensate for the lack of breastmilk-derived osteopontin. Whether the potential bioactivities are similar between intestinal-derived and breastmilk-derived osteopontin is not known.

We then sought to determine the effects of breastmilk-derived osteopontin on the development of the IEL compartment. Thus, we generated *Spp1*^+/-^ pups from littermate *Spp1*^+/-^ and *Spp1*^-/-^ dams crossed with *Spp1*^+/+^ sires and analyzed their IEL compartment at 3 and 8 weeks of life. Due to the great variation within samples in total IEL cellularity, our data are portrayed as cellular frequencies. At 3w, no differences were observed in TCRγδ^+^ and TCR^neg^ IEL (Figure 2 top panels). However, a significant reduction in frequencies of total TCRβ ^+^ IEL were observed in pups reared by osteopontin-deficient dams (Figure 2 middle panels). Within the TCRβ^+^ fraction, pups from *Spp1*^-/-^ dams had an increased proportion of CD4^-^CD8α^-^ IEL and a decreased proportion of CD4^+^ and CD4^+^CD8α ^+^ IEL (Figure 2, second row). The decrease in TCRβ^+^ IEL was also seen in the colon, and within this population, pups from *Spp1*^-/-^ dams had an increase in CD8αα^+^ IEL (Figure 3, bottom row).

**Figure 2:**
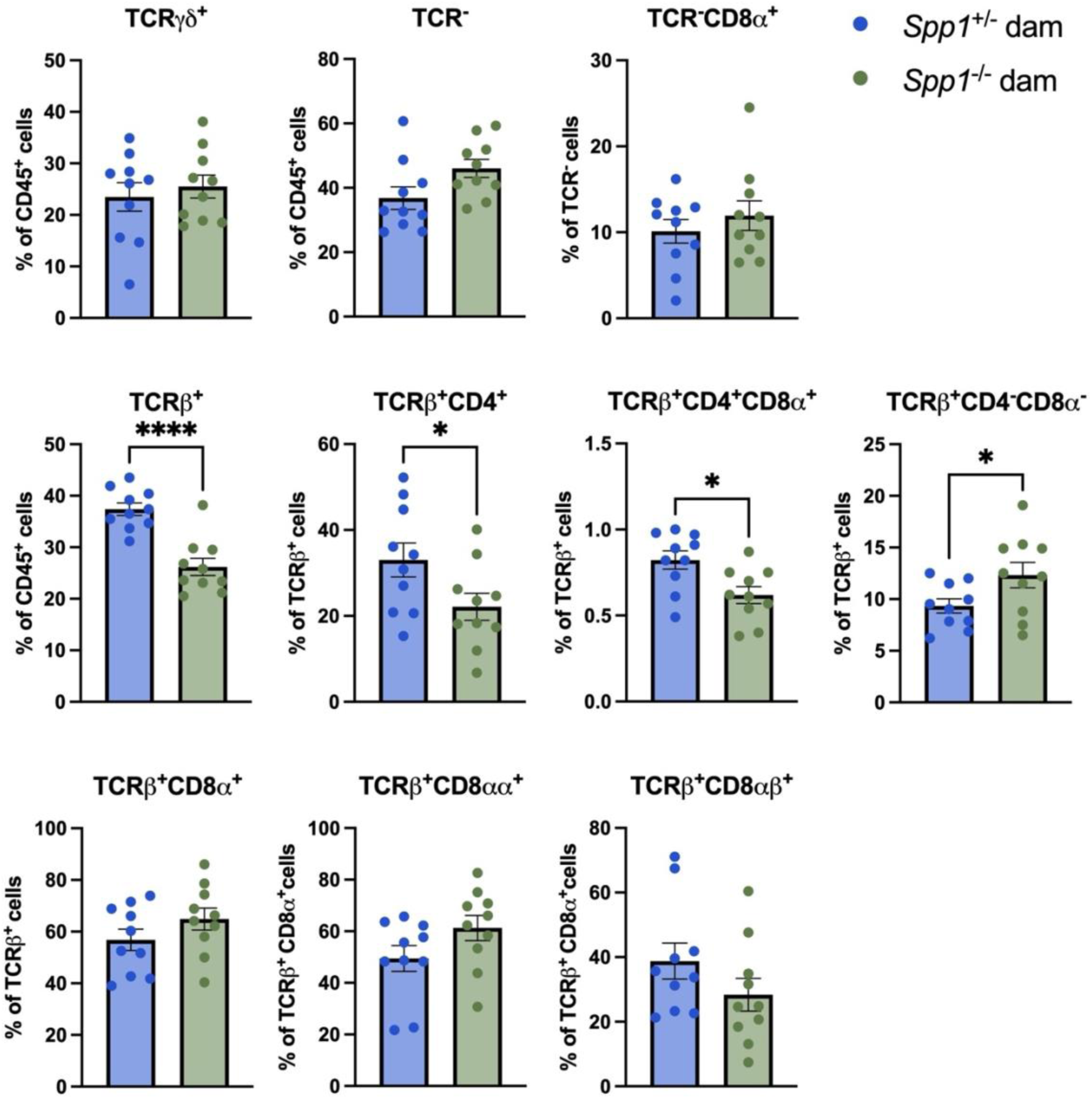
IEL frequencies in the small intestine of PND21 *Spp1*^+/-^ mice birthed and reared by *Spp1*^+/-^ and *Spp1*^-/-^ dams; two independent experiments; n=10. Student’s t-test, * p≤0.05 ****p≤0.0001.

**Figure 3:**
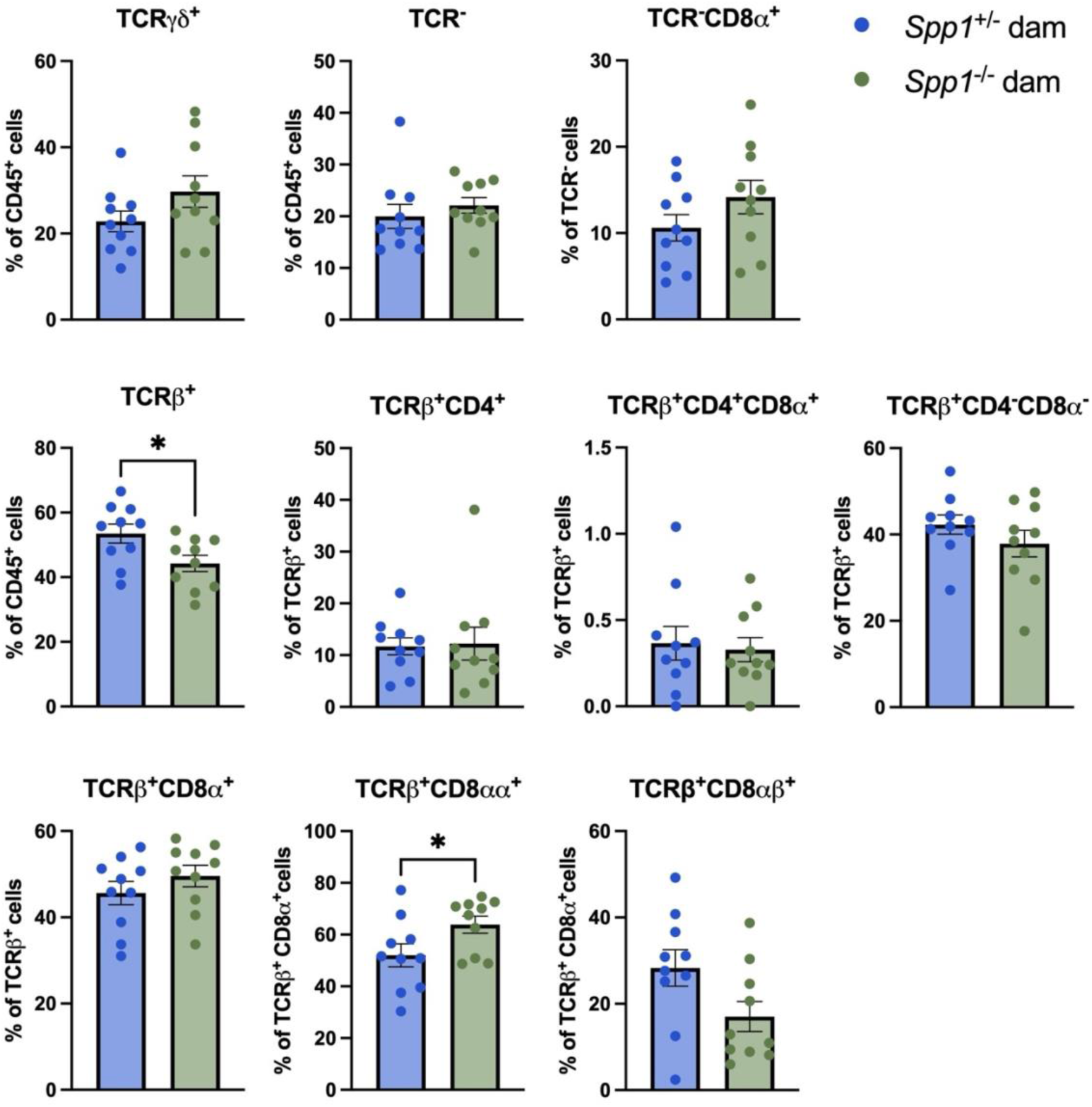
IEL frequencies in the colon of PND21 *Spp1*^+/-^ mice birthed and reared by *Spp1*^+/-^ and *Spp1*^-/-^ dams; two independent experiments. n=10. Student’s t-test, * p≤0.05.

At 8w, pups from *Spp1*^-/-^ dams had a decreased proportion of TCRγδ^+^ IEL. While the proportion of total TCRβ^+^ IEL was not different between groups, pups from *Spp1*^-/-^ dams had increased frequency of TCRβ^+^CD4^+^ IEL and a corresponding decrease in TCRβ^+^CD8α ^+^ IEL. Within the TCRβ^+^CD8α^+^ fraction, there was an observed decrease in CD8αα^+^ IEL and a corresponding increase in CD8αβ^+^ IEL (Figure 4, bottom panels). In the colon, pups from *Spp1*^-/-^ dams had an increase in TCR^neg^ IEL and TCR^neg^CD8α^+^ IEL. Within the TCRβ^+^ fraction, there was an increase in TCRβ^+^CD4^+^ IEL and a corresponding decrease in TCRβ^+^CD8α^+^ IEL (Figure 5), similar to that seen in the small intestine.

**Figure 4:**
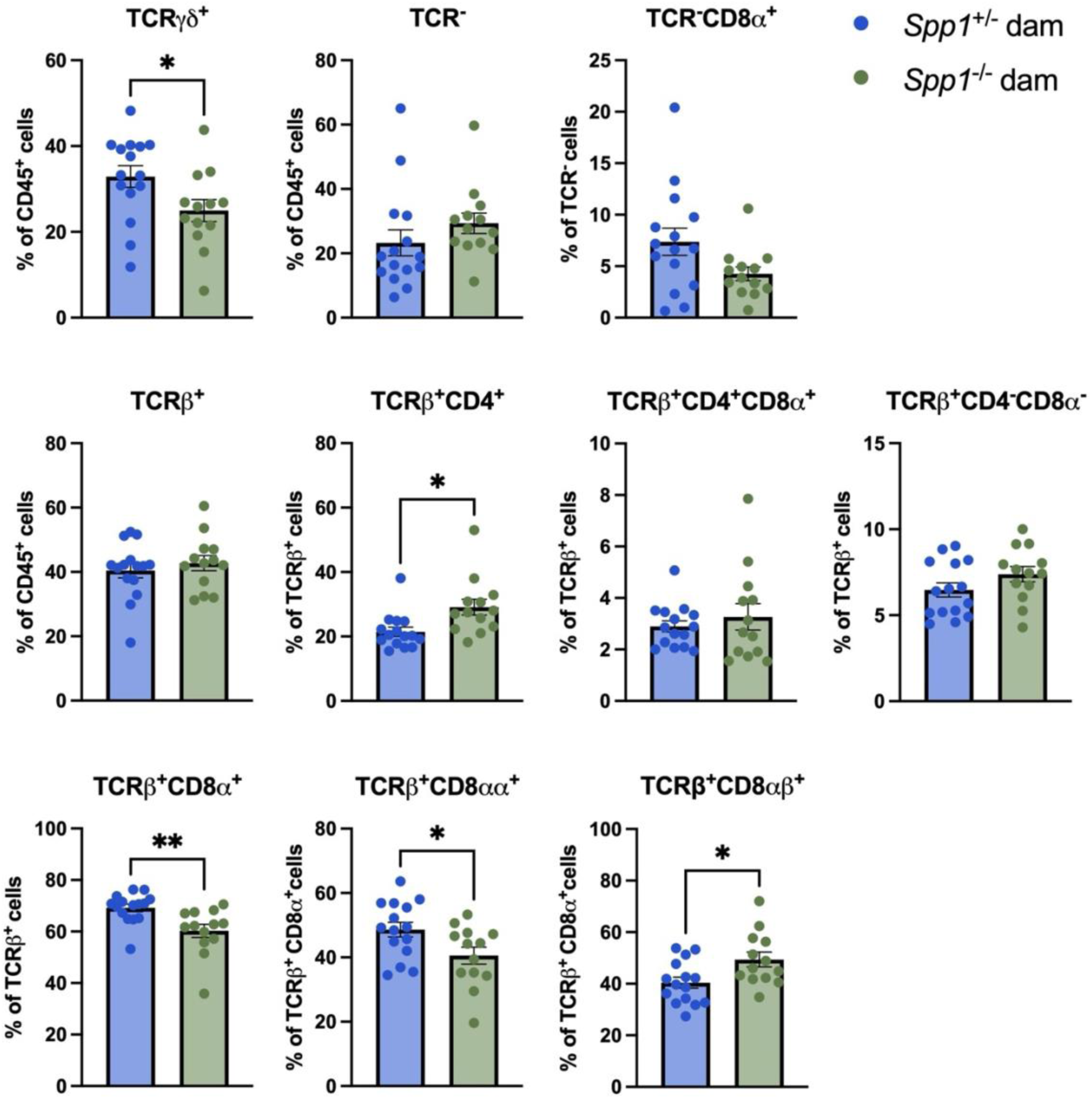
IEL frequencies in the small intestine of 8w *Spp1*^+/-^ mice birthed and reared by *Spp1*^+/-^ and *Spp1*^-/-^ dams; two independent experiments. n= 15 (*Spp1*^+/-^ dam), n=13 (*Spp1*^-/-^ dam). Student’s t-test, * p≤0.05, ** p≤0.01.

**Figure 5:**
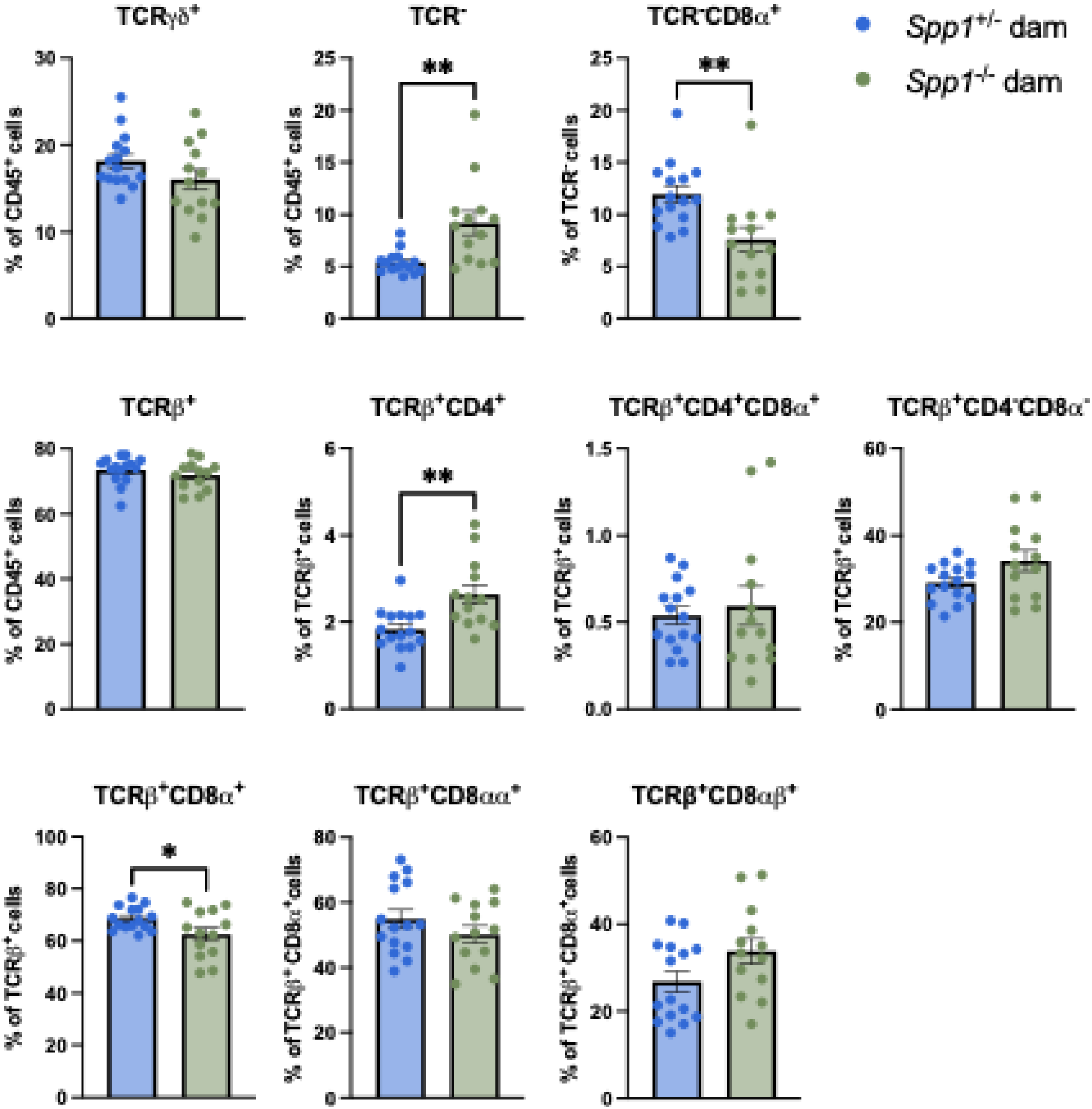
IEL frequencies in the colon of 8w *Spp1*^+/-^ mice birthed and reared by *Spp1*^+/-^ and *Spp1*^-/-^ dams; two independent experiments. n= 15 (*Spp1*^+/-^ dam), n=13 (*Spp1*^-/-^ dam). Student’s t-test, * p≤0.05, ** p≤0.01.

In summary, the IEL population frequencies in the small intestine and colon of pups derived from *Spp1*^+/-^ dams differed from pups reared by *Spp1*^-/-^ dams. The differences observed varied between recently weaned and adult mice, and also within specific IEL populations.

As this model was limited in pup genotype (only *Spp1*^+/-^ pups were analyzed) and only partially controlled for maternal microbiota effects, we generated a cross-fostered model in which littermate *Spp1*^+/-^ and *Spp1*^-/-^ dams were timed mated with *Spp1*^-/-^ sires. Upon birth of 3 synchronized litters (2 from *Spp1*^+/-^ dams and 1 from *Spp1*^-/-^ dam), dams were transferred between cages to foster each other’s pups (pups from *Spp1*^+/-^ dam to other *Spp1*^+/-^ dam, pups from *Spp1*^+/-^ dam to *Spp1*^-/-^ dam, pups from *Spp1*^-/-^ dam not used for this experiment (Figure 6).) This model controls for vertical transfer of microbiota during birth, as all pups analyzed in these experiments come from *Spp1*^+/-^ dams. Furthermore, this strategy generated both *Spp1*^+/-^ and *Spp1*^-/-^ pups, allowing for analysis of both maternal and pup contributions to intestinal osteopontin.

**Figure 6:**
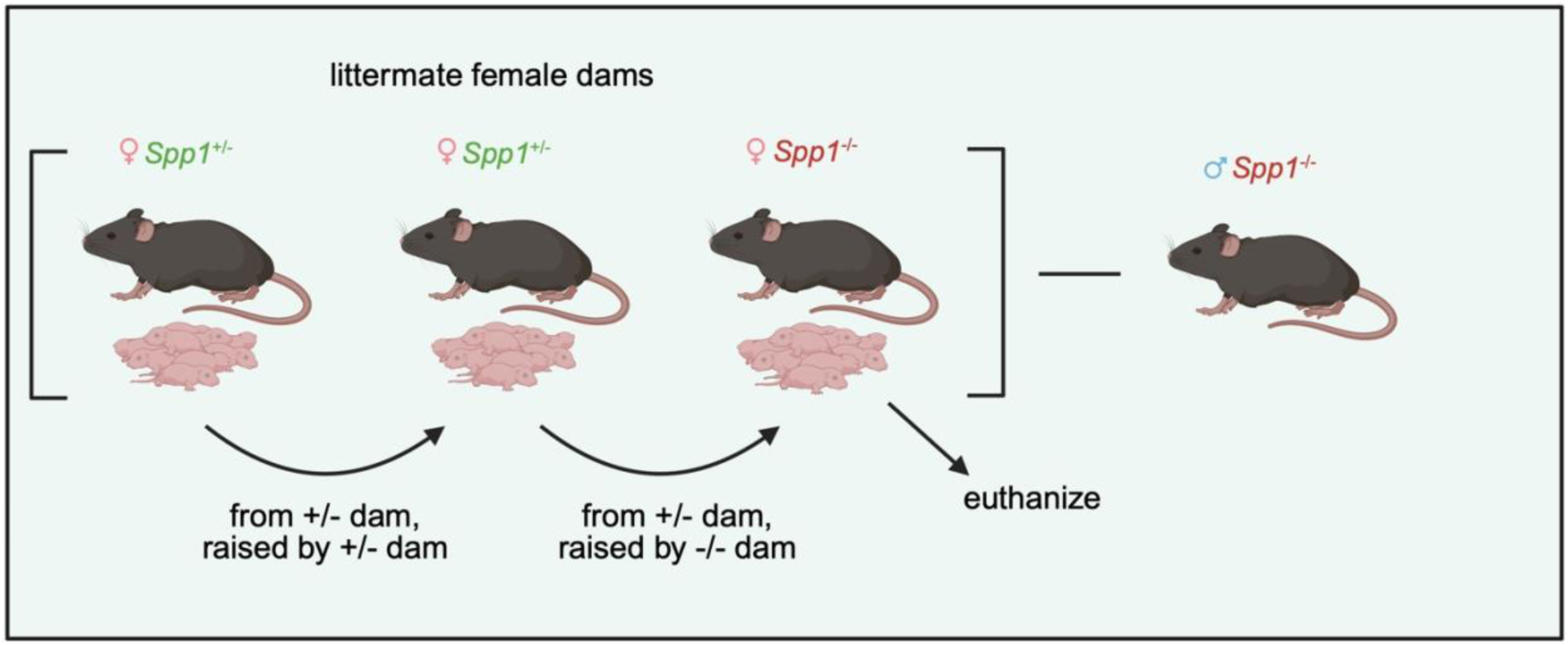
Breeding strategy for cross-fostered *Spp1* mice.

To determine the long-term effect of fostering with osteopontin-competent and -deficient dams, we analyzed the IEL compartment at 8w of age. To highlight the effect of maternal osteopontin, *Spp1*^+/-^ and *Spp1*^-/-^ mice from each dam are presented together. Complete data sets are available upon request. At 8w, mice fostered by osteopontin-deficient dams had a higher percentage of TCR^neg^ IEL, along with a lower percentage of TCRγδ^+^ and TCR^neg^CD8α^+^ IEL (Figure 7, upper panel). Within the TCRβ^+^ subset, mice fostered by osteopontin-deficient dams had a decreased percentage of CD4^+^CD8α^+^ IEL (Figure 7, middle panels) and an increased percentage of CD8αβ^+^ IEL.

**Figure 7:**
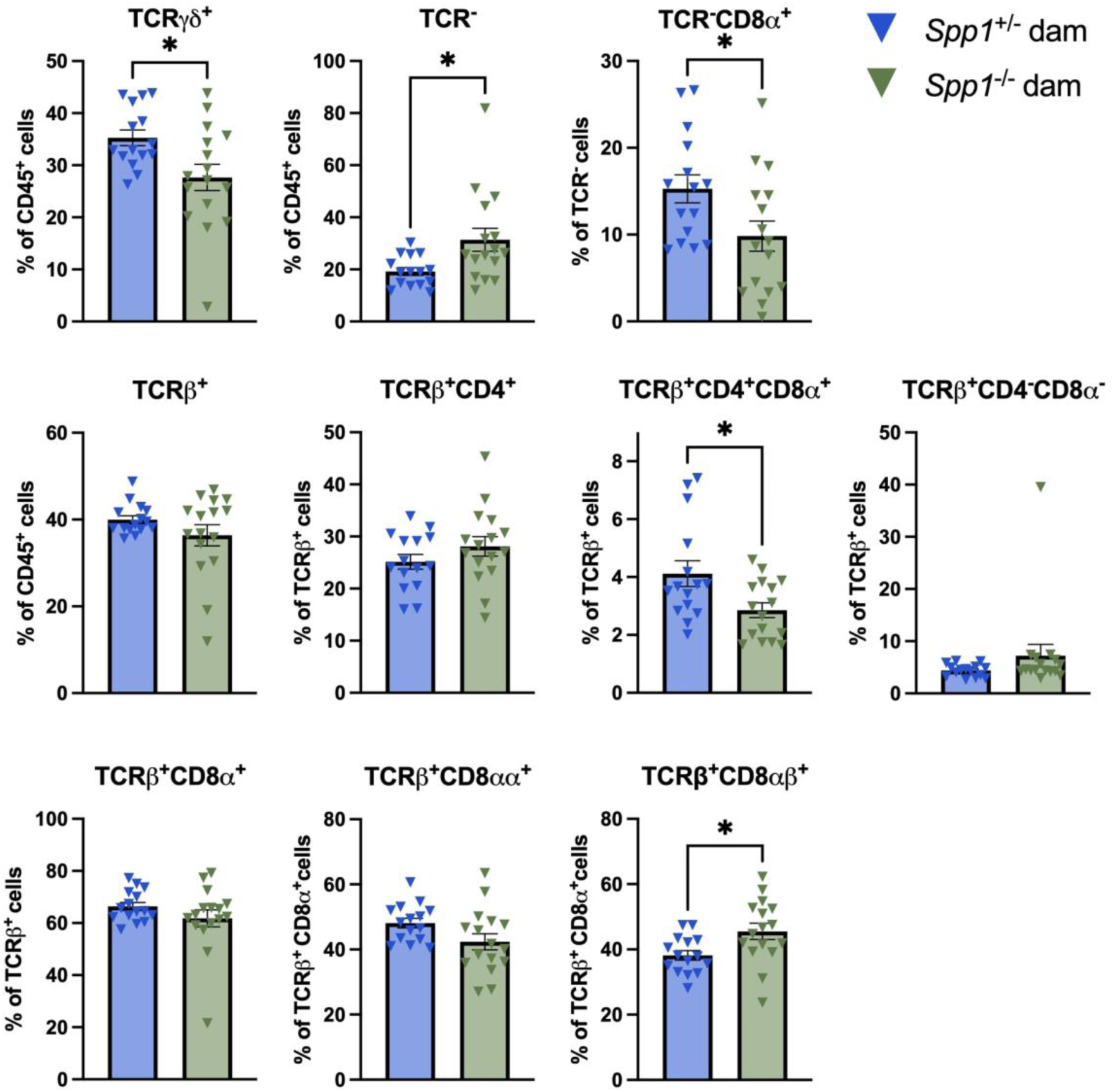
IEL frequencies in the small intestine of cross-fostered 8w mice; three independent experiments. n= 15 (*Spp1*^+/-^ dam), n=16 (*Spp1*^-/-^ dam). Student’s t-test, * p≤0.05.

These changes were less striking in the colon, though several changes were conserved between organs, most notably the decrease in TCRβ^+^CD4^+^CD8α^+^ IEL and the shift in CD8αα^+^/CD8αβ^+^ IEL toward increased CD8αβ^+^ cells (Figure 8).

**Figure 8:**
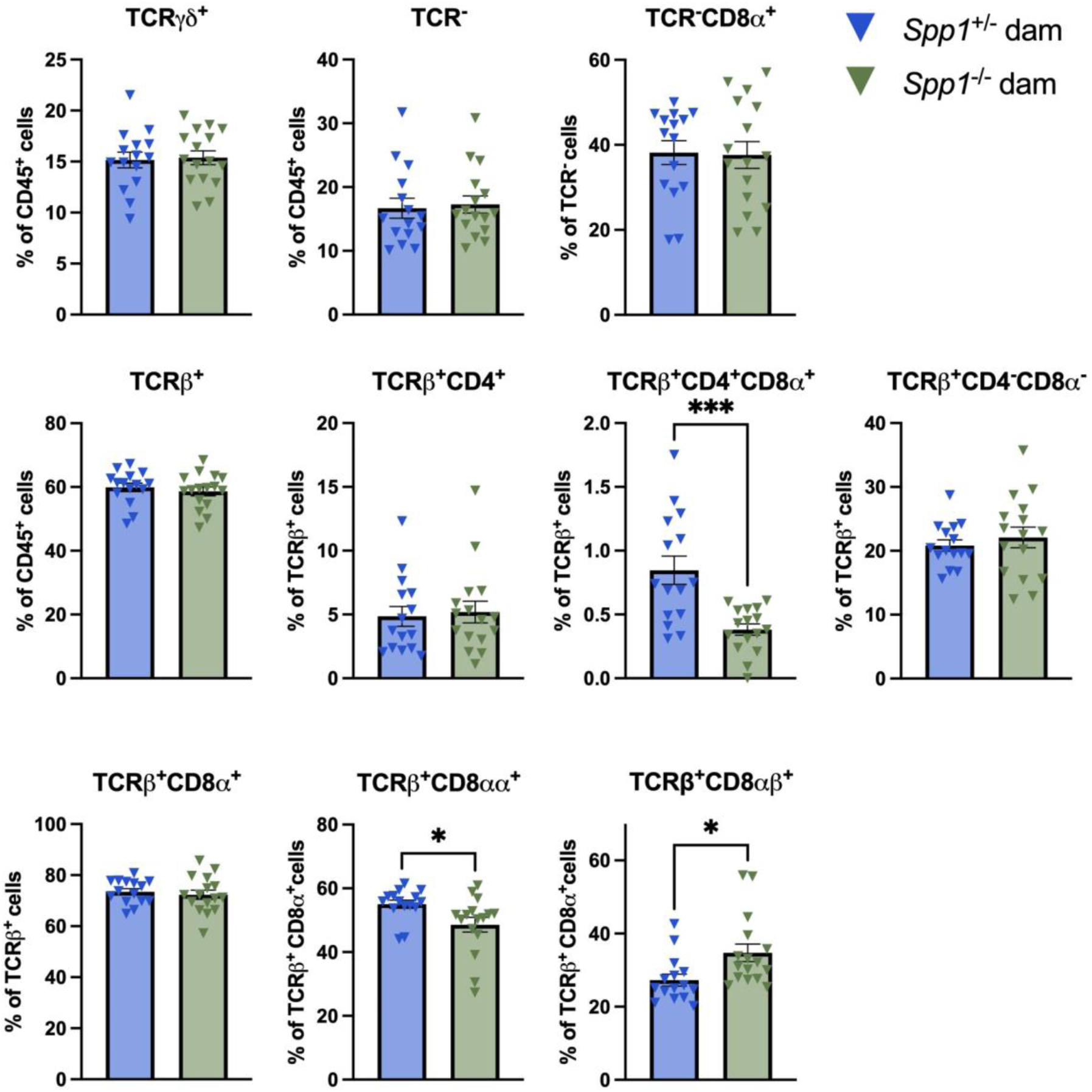
IEL frequencies in the colon of cross-fostered 8w mice; three independent experiments. n= 15 (*Spp1*^+/-^ dam), n=16 (*Spp1*^-/-^ dam). Student’s t-test, * p≤0.05, *** p≤0.001.

We reasoned that the observed differences in IEL frequencies in mice fostered by osteopontin-competent and -deficient dams may be a reflection of increased proliferation and/or cell death. However, Ki67 and annexin V analysis showed that the differences in IEL frequencies may not be associated with varied proliferation and cellular death between the two groups (Supplemental Figures 2 and 3).

We also analyzed the effect of maternal osteopontin contribution in peripheral sites by analyzing cellular composition of both the spleen and MsLN. Gross cellularity of the MsLN was similar between the groups, while in the spleen, mice fostered by osteopontin-deficient dams presented with increased percentage of CD19^+^ cells and a decrease in TCRβ^+^ cells, specifically TCRβ^+^CD4^+^ cells (Figure 9). Although significant, these differences were minor.

**Figure 9:**
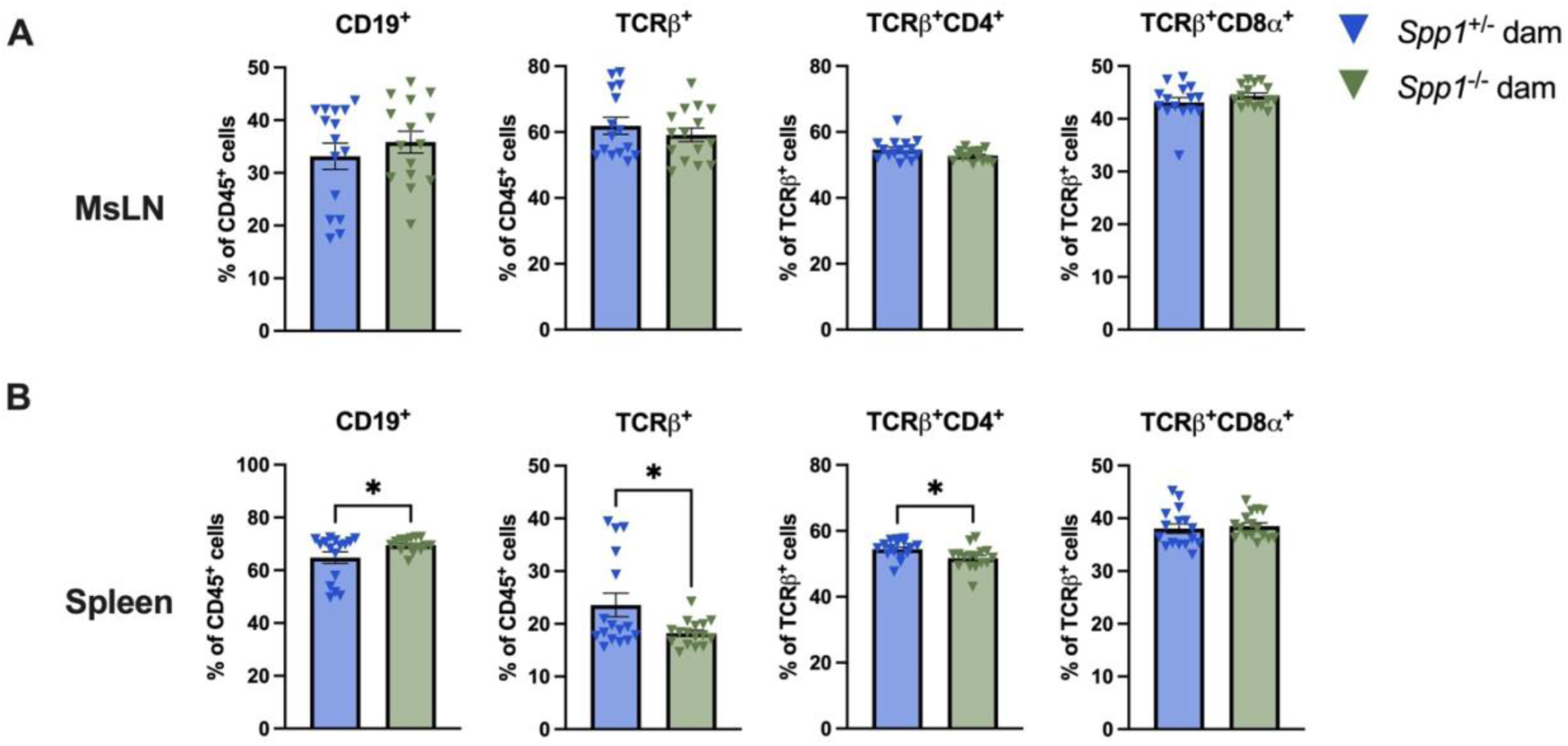
Frequencies of immune cells in lymphoid organs of 8w cross-fostered mice; three independent experiments; n=15 (spleen), n=16 (MsLN). Student’s t-test, * p≤0.05.

The diversity of IEL populations is accompanied with varied effector functions. However, because many IEL express granzymes (31, 32), it is believed that a great proportion of IEL are cytotoxic. To determine whether this particular effector function varied between IEL from 8w old mice fostered by *Spp1*^+/-^ and *Spp1*^-/-^ dams, we analyzed markers associated with cytotoxic function. While there were no differences in intracellular granzyme B and IFNγ levels between the two groups, a significant decrease in CD107a (a surrogate marker for degranulation) was observed int TCR^neg^ and TCRβ^+^ IEL in the small intestine and colon of mice fostered by osteopontin-deficient dams (Supplemental Figure 2). CD314 (a marker for cellular activation) was increased in TCRγδ^+^ IEL derived from mice fostered by osteopontin-deficient dams. These results suggest a potential dysfunction in the indicated IEL in mice raised by dams without breastmilk-derived osteopontin.

Due to relevant IEL function in intestinal homeostasis and inflammation (33–36), we investigated whether the observed changes in the IEL compartment in the intestines of mice fostered by osteopontin-deficient dams resulted in increased susceptibility to intestinal inflammation. We utilized several models of intestinal inflammation, including DSS-induced colitis (acute inflammation and recovery) and *Citrobacter rodentium* infection. No differences between 8w mice from osteopontin-sufficient or -deficient mice were observed (Figure 10).

**Figure 10:**
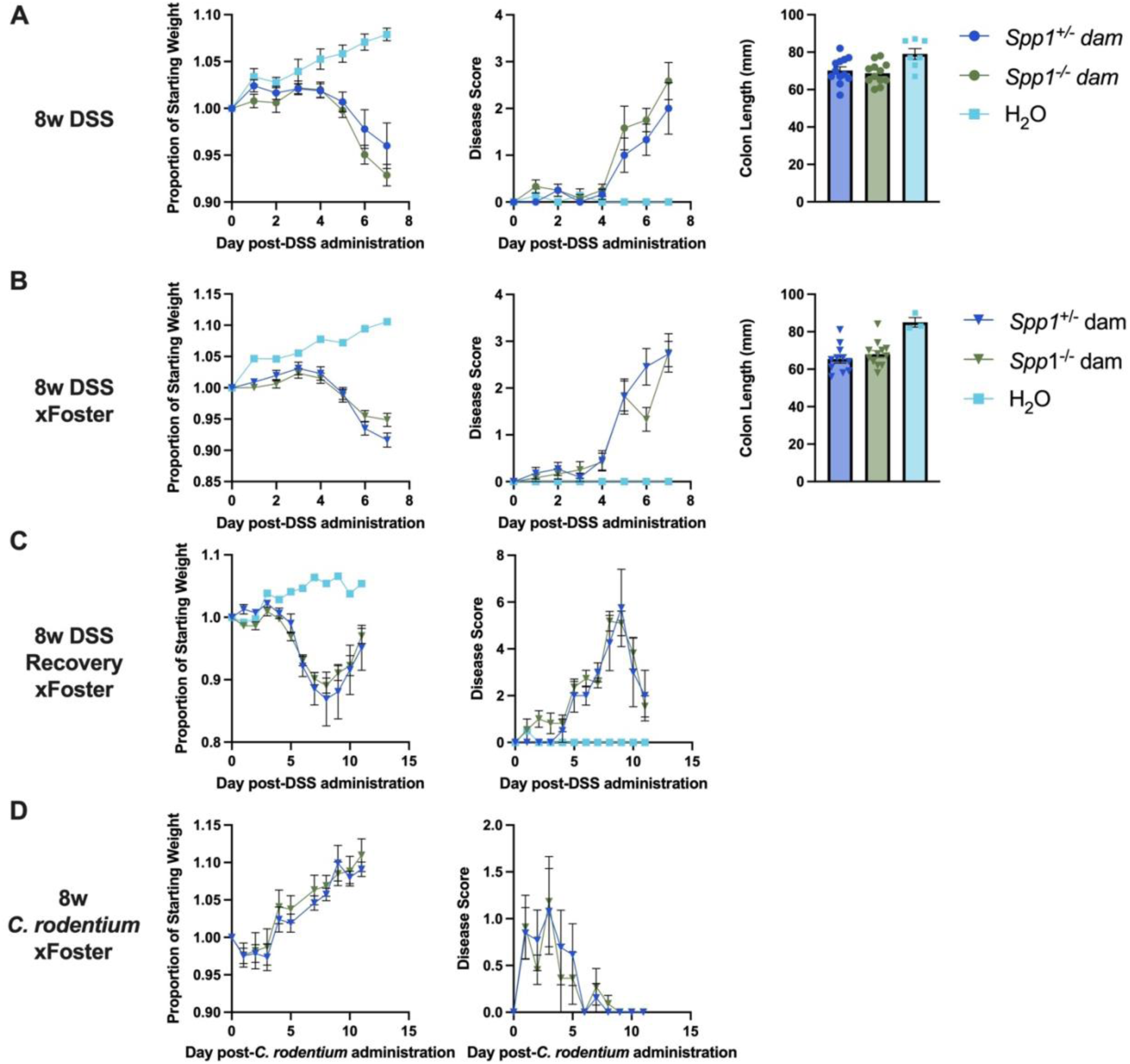
Response of breastmilk osteopontin-exposed and -non-exposed mice to models of intestinal inflammation. A, weight change (left), disease score (middle) and colon length (right) in the acute DSS model using *Spp1*^+/-^ mice birthed and reared by *Spp1*^+/-^ and *Spp1*^-/-^ dams; two independent experiments, n=12, 12, 7. B, weight change (left), disease score (middle) and colon length (right) in the acute DSS model using cross-fostered mice; one independent experiment, n=11, 12, 3. C, weight change (left) and disease score in the recovery DSS model with cross-fostered mice; three independent experiments, n=4, 11, 2. D, *Citrobacter rodentium* infection with cross-fostered mice; one independent experiment, n=13, 11.

## Discussion

In this report we show that the impact of breastmilk osteopontin on developing IEL is complex and multifaceted. Osteopontin affects different IEL subsets at different timepoints and induces a number of changes in these cells. Nevertheless, some conserved patterns were observed. In both mouse models, a shift in the proportion of small intestinal TCRβ^+^CD8αα^+^/TCRβ^+^CD8αβ^+^ IEL was observed, with mice from osteopontin-deficient dams displaying an increased proportion of TCRβ^+^CD8αβ^+^ IEL. Mice from these dams also displayed a decreased proportion of TCRγδ^+^ IEL in the small intestine. As >80% of TCRγδ^+^ IEL express CD8αα, this may indicate an effect of breastmilk osteopontin on cells expressing CD8αα. Adding further evidence to this theory is the observed effect on TCR^-^CD8α^+^, or iCD8α, cells. Alterations in this subset were seen in both mouse models in both the small intestine and colon, though not consistently in every experiment. Taken together, these data indicate that breastmilk osteopontin exposure has an effect on the development or stability of CD8αα^+^ IEL. The mechanism of this effect is beyond the scope of this publication and will require detailed and rigorous studies, likely at earlier time points, to investigate.

Although clear effects on the developing IEL compartment were observed in this study, the long-term effects of these changes are unclear. Despite differences in IEL composition, mice raised by osteopontin-deficient dams did not display increased susceptibility to DSS-induced colitis or to infection by *Citrobacter rodentium* at 8w of age. As the effects of early life osteopontin exposure appear to strongly affect cells expressing CD8αα, it is possible that a model more dependent on these cells would be a better option to further probe the effects of maternal osteopontin exposure. However, CD8αα+ IEL remain poorly understood, and their contributions to many intestinal diseases are unknown. It is also possible that the effects of maternal osteopontin exposure diminish over time, especially in osteopontin-competent mice. Exposing mice to models of inflammation and infection earlier in life (before 3w of age) may reveal more drastic phenotypes due to differences in the total amount of osteopontin present in the intestines at these times.

Differences between endogenous intestinal osteopontin and milk-derived osteopontin are poorly understood. We describe an effect based on maternal osteopontin exposure even when controlling for pup genotype (Figures 2-5), indicating that maternal osteopontin has a distinct function from the endogenous osteopontin found in the juvenile intestine. This may be partly due to an increase in total osteopontin concentration but is likely also due to differences between the isoforms. Milk osteopontin undergoes extensive post-translational modification (37) which may result in differences in bioactivity compared to endogenous intestinal forms. Milk osteopontin also passes through the stomach before arriving in the intestine, resulting in partial digestion and exposure of bioactive sites (38) In addition, milk-derived osteopontin has been shown to complex with lactoferrin, altering its bioactivity (39). The observed effects of milk-derived osteopontin likely result from the summation of several structural and functional differences.

The authors find it important to note that the *Spp1* cross-foster mice were not the only mice generated for this study. We began by generating a MMTV-Cre^+^ *Spp1*^fl/fl^ mouse by backcrossing mixed-background MMTV-Cre^+/-^ mice (Tg(MMTV-cre)4Mam/J, Jackson Labs #003553) onto our previously described *Spp1*^fl/fl^ mice (C57BL/6J-Spp1^em1Ddov^/J, Jackson Labs #037922) (40). These mice carried Cre recombinase under the control of the Mouse Mammary Tumor Virus Long Terminal Repeat promoter, which should have allowed for excision of the floxed allele more specifically to the mammary gland. However, these mice excised a floxed allele multiple times throughout their lineage, even in mice lacking expression of Cre recombinase. After intense screening and careful management, we were unable to ever recover these mice back to their *Spp1*^fl/fl^ background, which remains an unexplained genetic phenomenon. We have never seen this phenomenon in *Spp1*^fl/fl^ mice carrying Cre recombinase under any other promoter. The authors offer this as a note of caution to anyone attempting to generate these mice and suggest the use of a more tractable Cre line, such as the whey acidic protein Cre (WAP-Cre).

In summary, this report offers a preliminary description of the impact of breastmilk-derived osteopontin on the structure of the intestinal IEL compartment. Although the effects are varied, there is evidence that the most affected populations are those expressing CD8αα. Further studies will be needed to investigate the mechanisms by which breastmilk-derived osteopontin exerts its effect on IELs early in life, and the potential modulation of IEL effector functions.

## Supporting information

Table I. Antibodies used for staining

## Acknowledgements

Flow Cytometry experiments were performed in the VUMC Flow Cytometry Shared Resource. The VMC Flow Cytometry Shared Resource is supported by the Vanderbilt Ingram Cancer Center (P30 CA68485) and the Vanderbilt Digestive Disease Research Center (DK058404). Figure 6 was made using BioRender.

---

**Figure S1:**
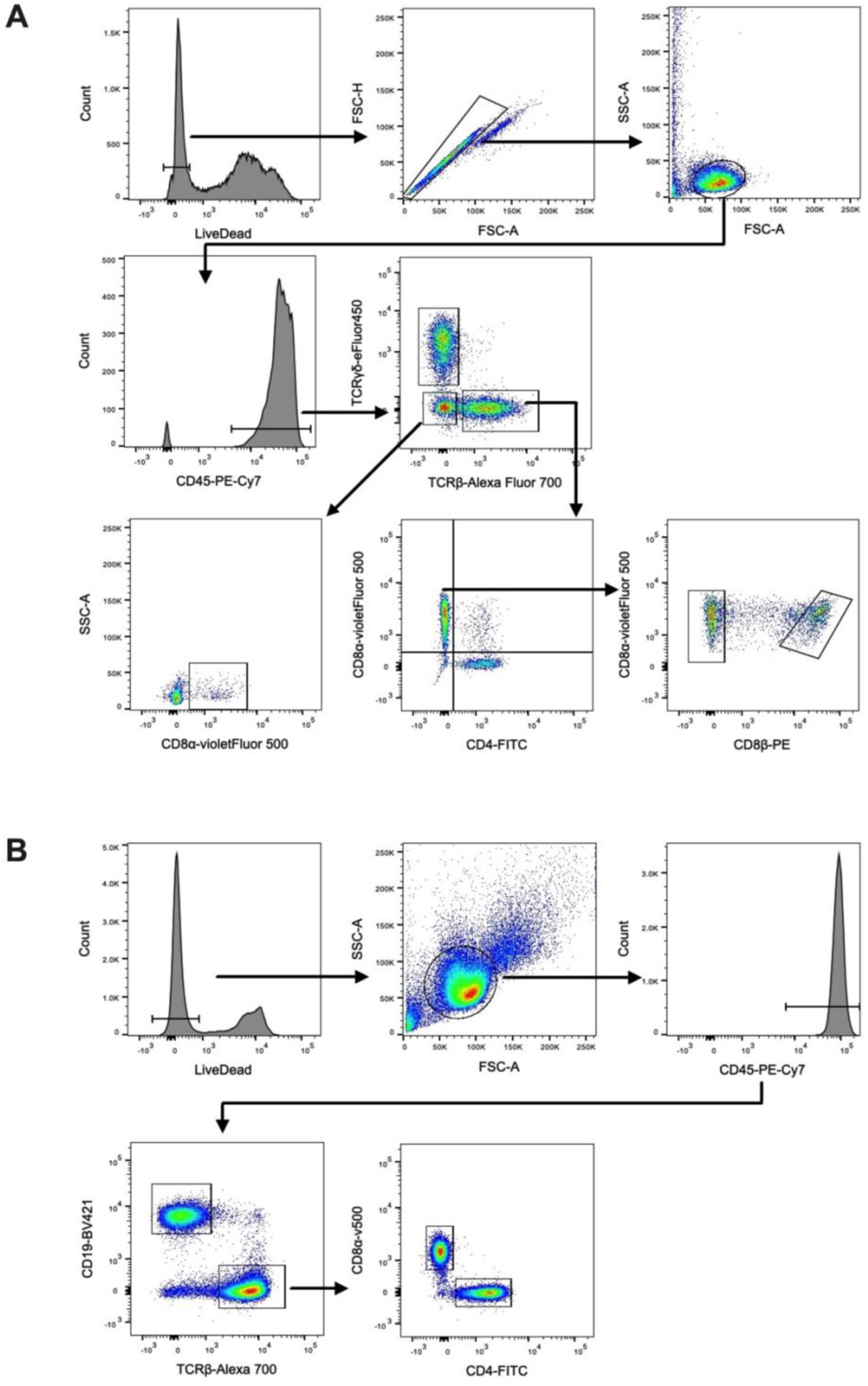
Gating strategies for A, IEL; and B, spleen and MsLN.

**Figure S2:**
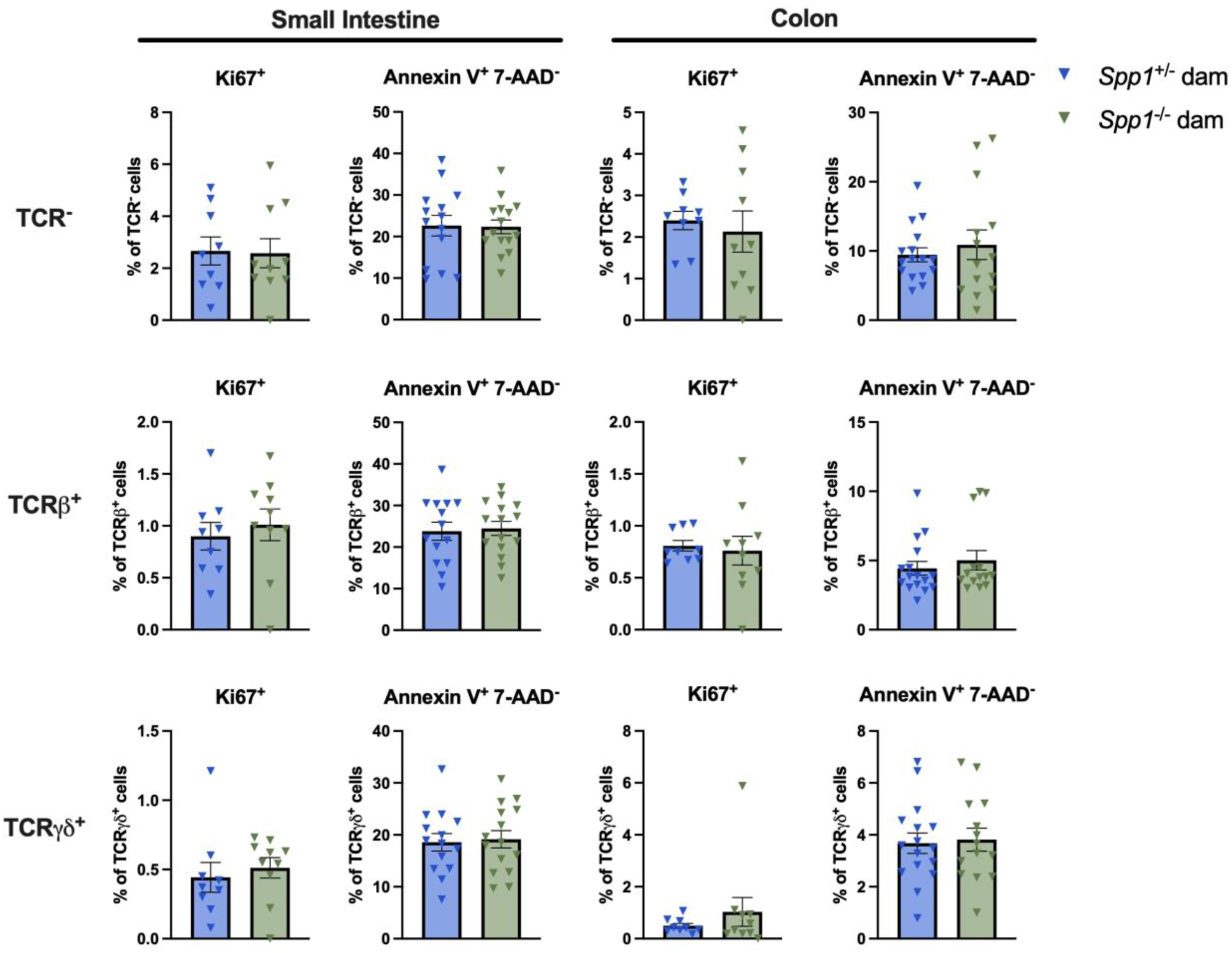
Cytotoxic and activation marker expression in IEL derived from 8w cross-fostered mice. A, small intestine; B, colon; three independent experiments. n= 15 (*Spp1*^+/-^ dam), n=16 (*Spp1*^-/-^ dam); Student’s t-test, * p≤0.05

**Figure S3:**
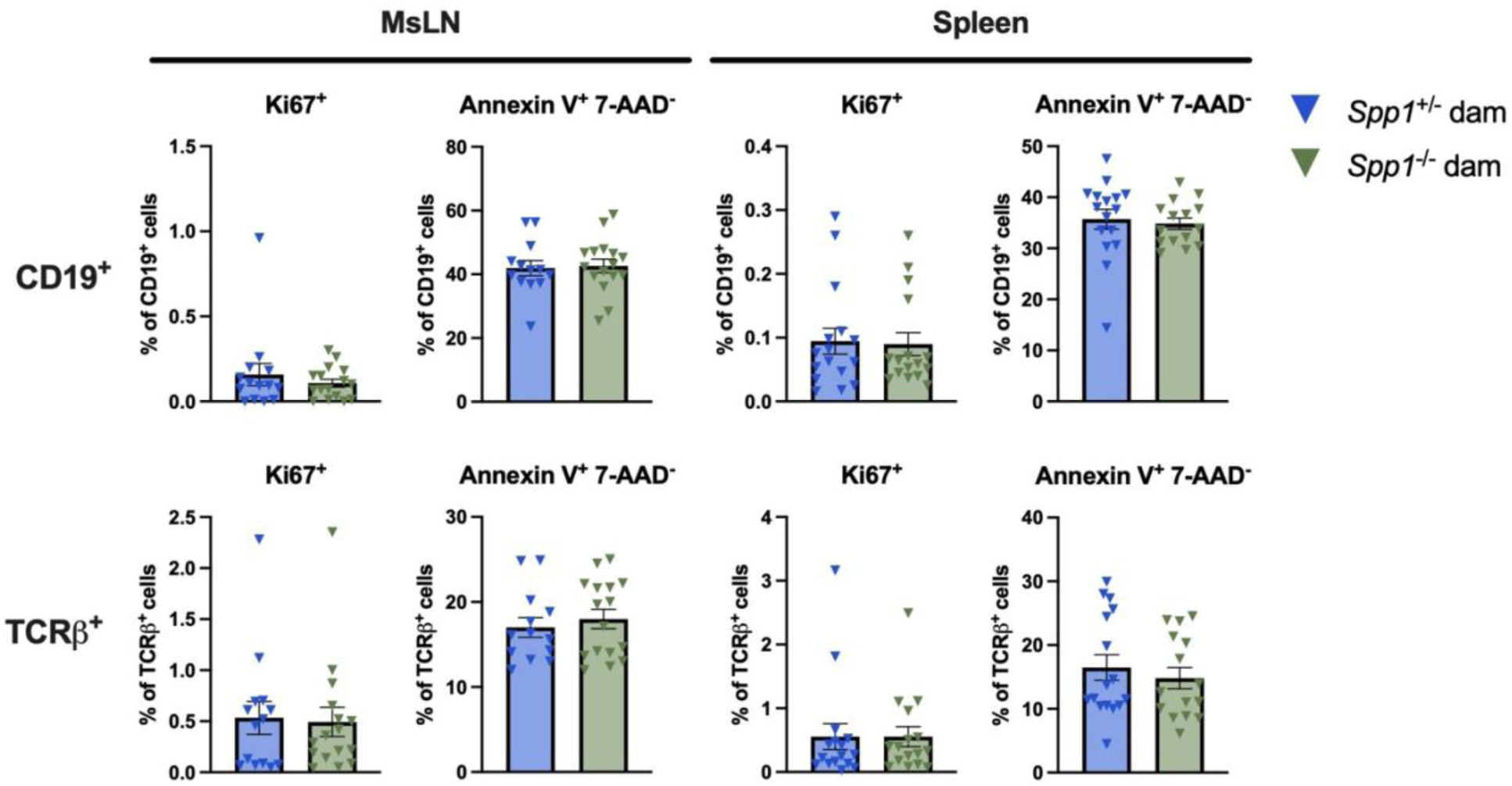
Ki67 and Annexin V staining of 8w cross-fostered IEL; three independent experiments. Ki67: n=9 (*Spp1*^+/-^ dam), n=10 (*Spp1*^-/-^ dam); Annexin V: n=14 (*Spp1*^+/-^ dam), n=15 (*Spp1*^-/-^ dam).

**Figure S4:**
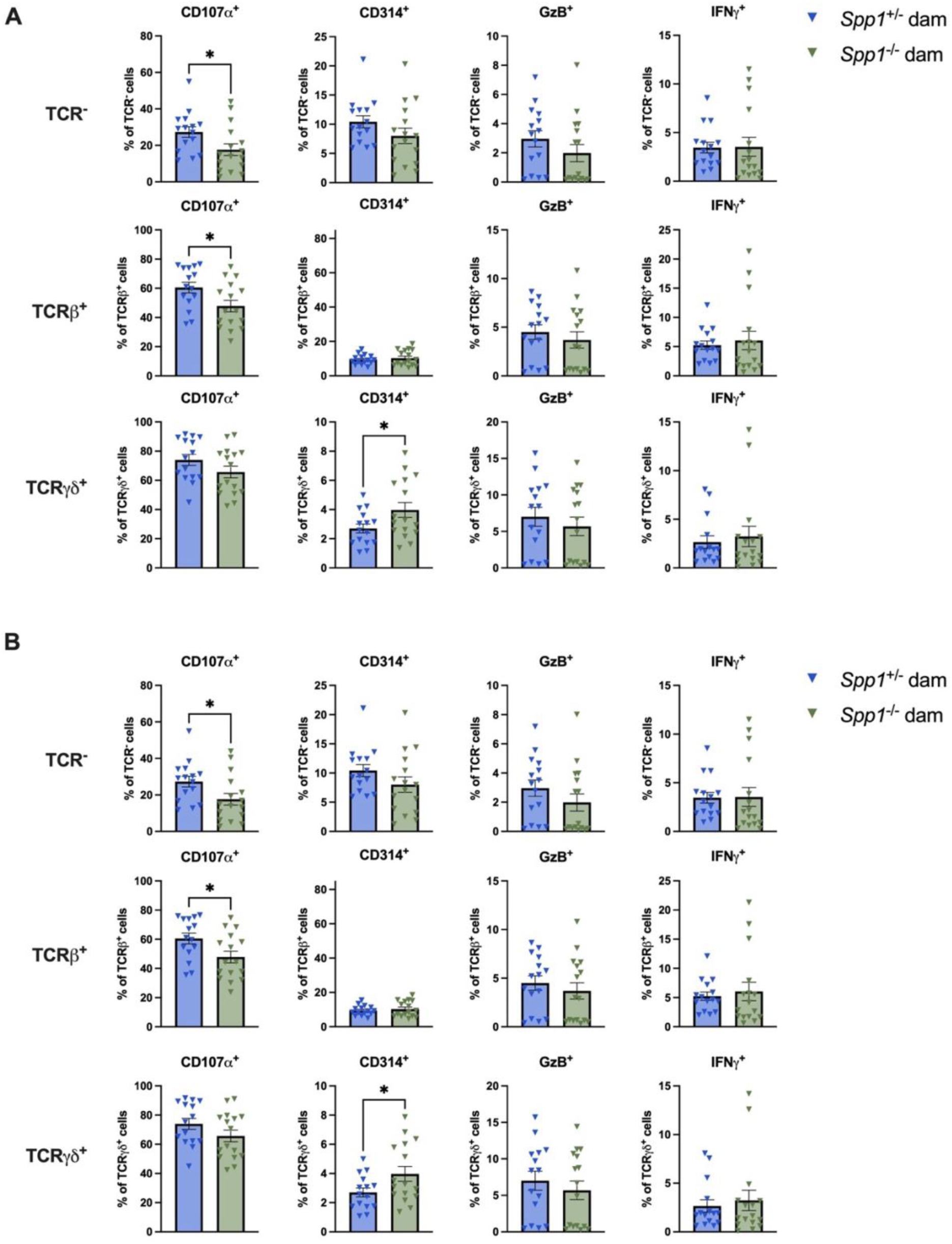
Ki67 and Annexin V staining of 8w cross-fostered secondary lymphoid organs; three independent experiments. MsLN Ki67: n=14 (*Spp1*^+/-^ dam), n=16 (*Spp1*^-/-^ dam); MsLN Annexin V: n=13 (*Spp1*^+/-^ dam), n=16 (*Spp1*^-/-^ dam); Spleen Ki67: n=16 (*Spp1*^+/-^ dam), n=16 (*Spp1*^-/-^ dam); Spleen Annexin V: n=16 (*Spp1*^+/-^ dam), n=15 (*Spp1*^-/-^ dam).

