## Supplementary material for "The effects of breastmilk-derived osteopontin on the intestinal intraepithelial lymphocyte compartment": Table I. Antibodies used for staining

| **Antibody** | **Clone** | **Supplier** | **Catalogue No.** | **Dilution** |
|---|---|---|---|---|
| Annexin V APC | - | Cytek Biosciences | 20-6409 | 1:100 |
| CD107a APC-Cy7 | 1D4B | BioLegend | 121615 | 1:200 |
| CD19 Biotin | 1D3 | Cytek Biosciences | 30-0193 | 1:200 |
| CD314 PE-Dazzle 594 | CX5 | BioLegend | 130213 | 1:200 |
| CD4 FITC | GK1.5 | Cytek Biosciences | 35-0041 | 1:200 |
| CD45 PE-Cy7 | 30-F11 | BD Biosciences | 561868 | 1:300 |
| CD8α APC-H7 | 53-6.7 | BD Biosciences | 560247 | 1:200 |
| CD8α violetFluor 500 | 53-6.7 | Cytek Biosciences | 85-0081 | 1:200 |
| CD8β PE | YTS156.7.7 | BioLegend | 126607 | 1:1000 |
| Ghost Dye$^{TM}$ UV 450 | - | Cytek Biosciences | 13-0868 | 1:1000 |
| Ghost Dye$^{TM}$ Violet 510 | - | Cytek Biosciences | 13-0870 | 1:1000 |
| GzB APC | NGZB | Invitrogen | 17-8898 | 1:200 |
| IFNγ PerCP-Cy5-5 | XMG1.2 | BioLegend | 505821 | 1:200 |
| Ki67 APC | 16A8 | BioLegend | 652405 | 1:200 |
| Streptavidin BV421 | - | BioLegend | 405226 | 1:200 |
| TCRβ Alexa Fluor 700 | H57-597 | BioLegend | 109223 | 1:200 |
| TCRβ PerCP-Cy5-5 | H57-597 | Cytek Biosciences | 65-5961 | 1:200 |
| TCRγδ eFluor450 | GL3 | Invitrogen | 48-5711 | 1:200 |

Table I – Antibodies used for extracellular and intracellular staining.
